# Electrophysiological Phenotype Characterization of Human iPSC-Derived Neuronal Cell Lines by Means of High-Density Microelectrode Arrays

**DOI:** 10.1101/2020.09.02.271403

**Authors:** Silvia Ronchi, Alessio Paolo Buccino, Gustavo Prack, Sreedhar Saseendran Kumar, Manuel Schröter, Michele Fiscella, Andreas Hierlemann

## Abstract

Recent advances in the field of cellular reprogramming have opened a route to study the fundamental mechanisms underlying common neurological disorders. High-density microelectrode-arrays (HD-MEAs) provide unprecedented means to study neuronal physiology at different scales, ranging from network through single-neuron to subcellular features. In this work, we used HD-MEAs *in vitro* to characterize and compare human induced-pluripotent-stem-cell (iPSC)-derived dopaminergic and motor neurons, including isogenic neuronal lines modeling Parkinson’s disease and amyotrophic lateral sclerosis. We established reproducible electrophysiological network, single-cell and subcellular metrics, which were used for phenotype characterization and drug testing. Metrics such as burst shapes and axonal velocity enabled the distinction of healthy and diseased neurons. The HD-MEA metrics could also be used to detect the effects of dosing the drug retigabine to human motor neurons. Finally, we showed that the ability to detect drug effects and the observed culture-to-culture variability critically depend on the number of available recording electrodes.

## 1. Introduction

The advent of efficient cellular reprogramming protocols has revolutionized stem cell research and enabled access to a large variety of human cell lines^1^. Induced-pluripotent-stemcell (iPSC)-technology has been used to establish 2D and 3D human cell and tissue models^2^ and to study organ genesis for, e.g., liver^3^, lungs^4^, heart^5^ and brain^6^. Today, iPSC technology allows to obtain cells and tissue samples from both, healthy individuals and patients and to develop *in-vitro* disease models^7^. Genetically-defined human iPSC-derived neurons have been generated to investigate physiological aspects and mechanisms underlying common neurological disorders^8^ and for usage in *in-vitro* platforms for systematic drug testing^9,10^. Functional phenotype characterization and drug-effect assessment of human neurons can be performed *in vitro* by using optical methods (e.g., calcium imaging^11^), patch-clamp techniques^12^, or by using microelectrode arrays (MEAs)^13^. MEAs feature a set of metal electrodes to record electrical activity simultaneously from several hundreds to thousands of cells and have been used to study brain disorders and to detect differences between normal and pathological conditions^14,15^.

Passive MEA devices^16–18^ without active circuit elements feature 10-100 electrodes per mm^2^ and have been used to electrically characterize human neuronal phenotypes^9,10,14^. Complementary-metal-oxide-semiconductor (CMOS)-based high-density microelectrode arrays (HD-MEAs)^19–24^ feature active circuit elements and electrode densities of several hundreds to thousands of electrodes per mm^2^ and have been used in first experiments to characterize electrical properties of healthy human iPSC-derived neurons^25^. However, to the best of our knowledge, CMOS-based HD-MEAs have not been yet used to comprehensively characterize and compare electrical phenotypes of human neuronal line across scales including network, single-neuron and subcellular features and to assess the effects of neurological disorders.

Among neurodegenerative disorders, Parkinson’s disease and amyotrophic lateral sclerosis have been extensively studied by using iPSC technology in order to understand basic disease mechanisms at molecular and functional levels and for finding new cures^26–30^.

Parkinson’s disease (PD) is among the most common neurodegenerative diseases, with a prevalence increasing from 2.5 million patients in 1990 to 6.1 million patients in 2016^31^, and projections indicating that more than 12 million people will be affected by 2050^31^. PD is characterized by the death of dopaminergic neurons within the *substantia nigra*, which causes progressive motor and cognitive dysfunctions, such as tremor, rigidity and dementia^32^. The distinctive hallmark of PD is the accumulation of intracellular α-synuclein, which forms protein inclusions, known as Lewy bodies and Lewy neurites^32^. Human iPSC-derived neurons modeling PD have been studied *in vitro* and showed compromised neuronal morphology^33^, synaptic connectivity^34^ and different electrophysiological characteristics including, e.g., the shape of action potential waveforms^35^.

Amyotrophic lateral sclerosis (ALS) is a neurodegenerative disease, characterized by a progressive degeneration of upper and lower motor neurons, followed by muscle degeneration, paralysis, and respiratory failure^36^. ALS incidence is reported to be between 0.6 and 3.8 per 100’000 persons/year^37^. Human iPSC-derived neurons, developed to model ALS, featured an intrinsic membrane hyperexcitability phenotype^10^, which was used to test compounds that lowered neurons’ electrical hyperexcitability. The study of Wainger et al. using 64-microelectrode arrays led to clinical testing of retigabine in human patients^10^. Combining human iPSC-derived neurons with large-scale electrophysiological techniques, such as HD-MEAs, provides a powerful and scalable platform to study neurological disease mechanisms and offers the potential to assess disease-induced phenotypic alterations.

Here, we used CMOS-based high-density microelectrode arrays (HD-MEAs) to characterize and compare the electrical phenotypes of four iPSC-derived neuronal cell lines: human dopaminergic neurons (hDNs), dopaminergic neurons carrying the A53T α-synuclein mutation, linked to PD (hDNs-PD), motor neurons (hMNs), and motor neurons carrying the TDP-43 Q331K mutation, linked to ALS (hMNs-ALS). We found significant functional phenotype differences across the studied iPSC lines at different spatiotemporal scales, ranging from network level to characteristics of individual axons. We then used HD-MEA-based readouts to quantify the effect of the drug retigabine on motor neurons (hMNs)^10,38^ and we showed that high-spatiotemporal-resolution sampling of neuronal activity by means of HD-MEAs provides very reproducible readouts in assessing the effects of neuroactive compounds.

## 2. Results

### 2.1. Human iPSC-derived neurons develop spontaneous activity on HD-MEA chips

Before probing the developing neurons and neuronal networks, we confirmed the presence of characteristic cell-type specific neuronal markers for each iPSC line. Human iPSC-derived and rat primary neuronal cultures, used for benchmarking, showed electrical activity across the entire HD-MEA chip electrode area (Figure 1a). The electrically active areas contained mature neurons as confirmed by microscope imaging of MAP2-positive cells (Figure 1b). The presence of astrocytes in the cell cultures was confirmed immunohistochemically by GFAP^39,40^ and S100-β^40^ staining (Figure 1c). Furthermore, we visualized motor and dopaminergic neurons on the HD-MEA chips using the motor-neuron marker SMI-32^39^ (Figure 1c, central panel) and the dopaminergic-neuron marker TH^41^ (Figure 1c, right panel). Human and rat neurons were all electrically active and showed spontaneous action-potential signals, which systematically crossed the detection threshold. (Figure 1d).

**Figure 1.**
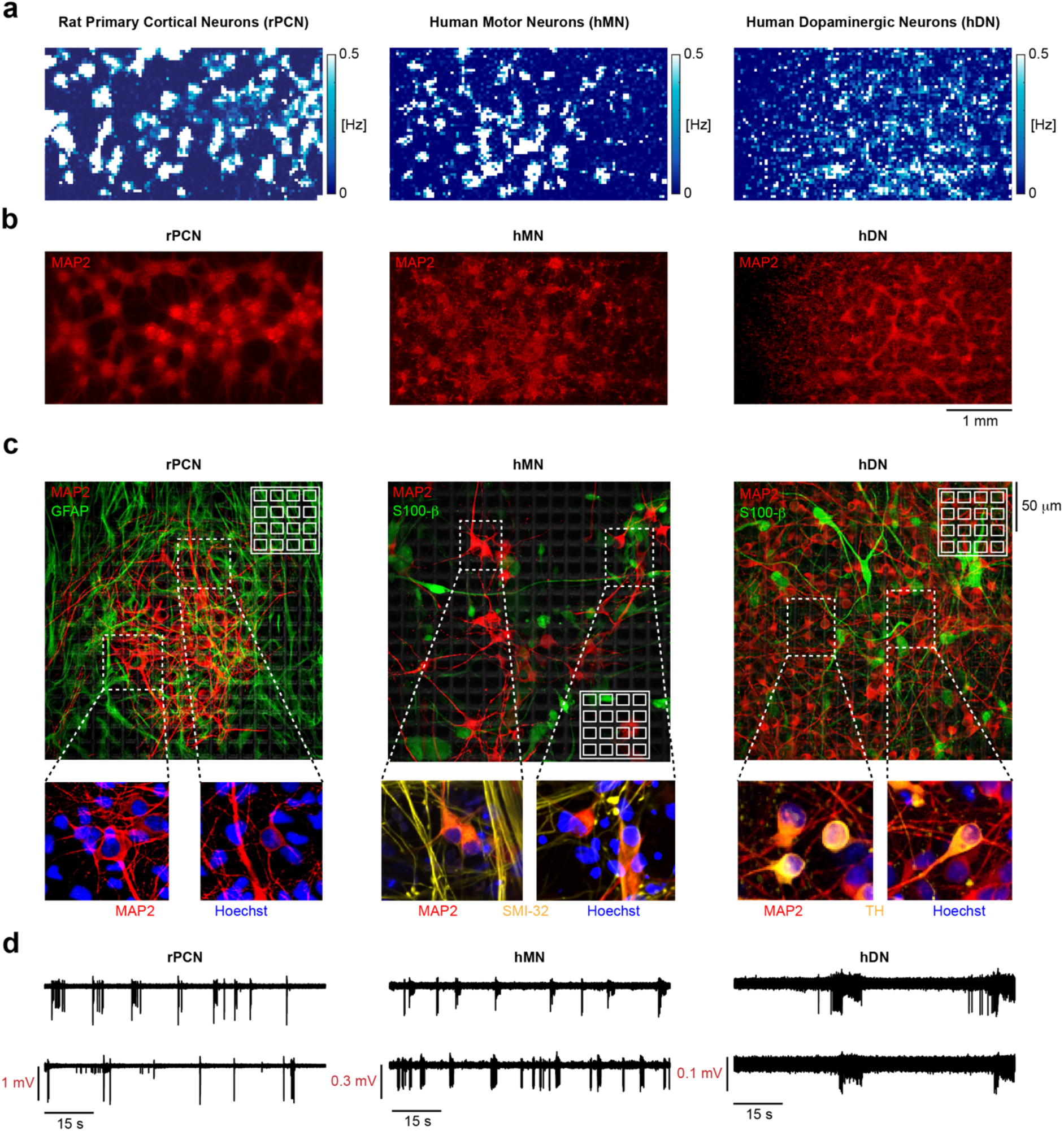
Electrical and optical imaging of neuronal cultures. **(a)** HD-MEA electrical image showing 2D spatial distribution maps of the electrode firing rate (eFR, see Methods, *HD-MEA Metrics)* as recorded across the entire HD-MEA chip surface from 6’600 electrodes at DIV 21, for rat primary cortical neurons (rPCNs), human motor neurons (hMNs) and human dopaminergic neurons (hDNs). **(b)** Microscope image of MAP-2 positive (red) stained neurons on HD-MEA chips. Cells were fixed and stained on the HD-MEA chips at DIV 21. **(c)** Cell-type specific stainings of cultures on HD-MEA chips shown in Figure 1b. Cell nuclei are shown in blue (Hoechst positive), motor neurons (SMI-32 positive) and dopaminergic neurons (TH positive) in yellow, astrocytes (GFAP, S100-β positive) in green. **(d)** Example voltage traces showing extracellular action potentials (spikes) recorded by two electrodes at DIV 21. Left panel: rat primary cortical neurons. Central panel: human motor neurons. Right panel: human dopaminergic neurons.

### 2.2. Electrical phenotype characterization of human iPSC-derived neurons across development

To compare the electrophysiological properties across the four human iPSC-derived neuronal cell lines, we first examined and compared the mean firing rate (MFR), mean spike amplitude (MSA), mean inter-spike interval (ISI) coefficient of variation (ISIcv) and the percentage of active electrodes (pAE). Changes in metrics as the firing rate are hallmarks of neuronal development^10^, but they can also be indicative of a specific neuron type^42^ or pathology^10^. We also compared the iPSC-derived cell lines electrophysiological properties with the ones of rat primary cortical neurons (rPCNs), the most commonly used neuronal cells in the microelectrode-array field and an established *in-vitro* culturing system, across development (for details, see Figure S1). Both, motor (hMN) and dopaminergic (hDN) neuronal lines showed a significant increase (7 fold and 1.9 fold) in the mean firing rate from DIV 7 to DIV 21 (Figure 2a, 2b), similar to rPCNs (Figure S1c).

**Figure 2.**
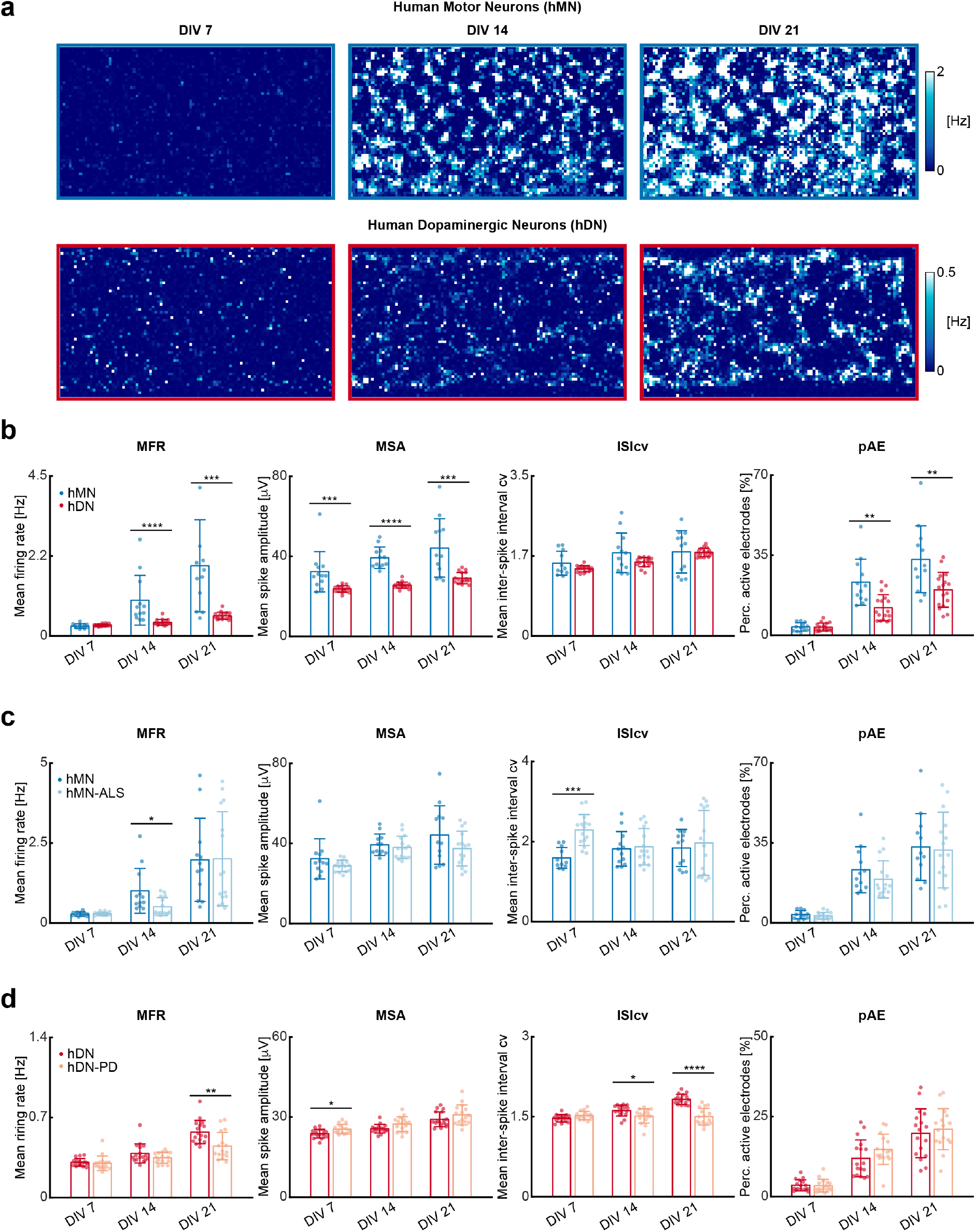
Electrical phenotype characterization of human iPSC-derived neurons across development. **(a)** Exemplary 2D spatial distribution maps (6’600 electrodes) of electrode firing rates for motor neurons (top) and dopaminergic neurons (bottom), at DIVs 7, 14 and 21, respectively. **(b)** Bar plots comparing the mean firing rate (MFR), mean spike amplitude (MSA), mean ISI coefficient of variation (ISIcv) and percentage of active electrodes (pAE) of 12 hMN cultures (blue) and 17 hDN cultures (red) at DIVs 7, 14 and 21. **(c)** Bar plots comparing the mean firing rate, mean spike amplitude, mean ISI coefficient of variation and percentage of active electrodes of 12 hMN cultures (blue) and 15 hMN-ALS cultures (light blue) at DIVs 7, 14 and 21. **(d)** Bar plots comparing the mean firing rate, mean spike amplitude, mean ISI coefficient of variation and percentage of active electrodes of 17 hDN cultures (red) and 16 hDN-PD cultures (orange) at DIVs 7, 14 and 21. Each dot represents one HD-MEA or well. Bar heights indicate distribution mean values, and error bars indicate standard deviations. The black stars indicate p values: * p < 0.05, ** p < 0.01, *** p < 0.001, **** p < 0.0001.

hMNs featured a higher mean firing rate, mean spike amplitude and percentage of active electrodes compared to hDNs (Figure 2b). In particular, at DIV 21, the hMN mean firing rate (1.98 ± 1.30 Hz) was 3.5 times higher (p < 0.001) than the hDN mean firing rate (0.57 ± 0.10 Hz). The hMN mean spike amplitude and percentage of active electrodes were 1.5-fold and 1.7-fold higher than that of the hDN line (Figure 2b). However, the mean ISI coefficient of variation did not vary significantly across development between hMNs and hDNs, which showed a similar behavior in neuron-firing-rate evolution over development.

Both, hMNs and hDNs showed significantly lower mean spike amplitudes in comparison to rPCNs at DIV 21. rPCNs had an average mean spike amplitude of 95.01 ± 8.96 μV, whereas hMNs and hDNs featured mean spike amplitudes of 44.23 ± 14.62 μV and 29.17 ± 2.78 μV, respectively (Figure S1c and Figure 2b). Furthermore the variation in the mean spike amplitude across development was significantly less (< 1.2 fold) in hMNs and hDNs than that of rPCNs (Figure S1c and Figure 2b). Human iPSC-derived neuronal lines were also characterized by an almost constant ISI coefficient of variation across development, in contrast to the increase in rPCNs (Figure S1c and Figure 2b).

Comparing healthy (WT) and diseased (ALS) motor neuron lines, the hMN-ALS neuronal cells featured a 0.5-fold (p < 0.05) lower mean firing rate than hMNs at DIV 14 (Figure 2c). Similar results were found by comparing the two dopaminergic neuron lines, which showed a 0.8-fold (p < 0.01) lower mean spiking rate of hDN-PDs in comparison to hDNs at DIV 21 (Figure 2d). Analysis of the mean spike amplitudes showed no significant differences between hMN and hMN-ALS neurons. In contrast, the two dopaminergic neuron lines, hDN and hDN-PD, showed significant differences in the mean spike amplitude at DIV 7 (p < 0.05) (Figure 2c).

The mean ISI coefficient of variation evidenced a more regular firing of the hMN line at DIV 7 in comparison to the hMN-ALS line (Figure 2c). Conversely, the disease line hDN-PD featured a more regular firing at DIV 21 in comparison to the healthy hDN line (Figure 2d). Finally, by comparing the percentage of active electrodes across development, we did not find significant differences between hMN and hMN-ALS or hDN and hDN-PD lines, suggesting that potential differences in electrical phenotypes are likely not a consequence of differences in the number of active cells between healthy and diseased lines. (Figure 2c and 2d).

### 2.3. Network burst characterization of human iPSC-derived neurons across development

Neuronal networks are often characterized by synchronous activity (bursts), which may give rise to network oscillations^43^. These bursts emerge as a result of recurrent synaptic connections that form as neuronal networks mature. In primary cortical neurons, the bursts are often irregular even in mature networks (Figure S2). The nature and properties of the associated oscillations may vary for different pathological conditions ^44^ or in co-cultures with different cell types^42^.

Here, we assessed the characteristic properties of human healthy and diseased cell lines by using network-burst metrics including mean burst duration (BD), mean inter-burst interval (IBI), mean IBI coefficient of variation (IBIcv) and burst shape (see Methods, *HD-MEA Metrics*).

All human iPSC-derived neuronal cultures showed robust and reproducible network oscillations at DIV 14 and 21 (Figure 3a, Figure S2). hMNs and hDNs featured significant differences in IBI and IBIcv at both DIVs 14 and 21 (Figure 3b). hDNs featured a 2.4-fold increase (p < 0.0001) in time between consecutive bursts compared to hMNs at DIV 21 (Figure 3b). Furthermore, hDNs displayed more regular bursts than hMNs, which is shown by a 0.3-fold decrease (p < 0.001) in IBIcv between consecutive bursts at DIV 21 (Figure 3b). Comparing motor-neuron lines, we found that hMN-ALS neurons featured a longer burst duration of 16.56 ± 5.44 s than that of hMNs (3.91 ± 2.59 s) at DIV 21 (Figure 3c). Converse results were found for the two isogenic dopaminergic-neuron lines, where hDN-PDs showed a 0.5-fold decrease (p < 0.0001) of the burst duration at DIV 21 compared to hDNs (Figure 3d). By comparing the inter-burst interval times of healthy and diseased lines at DIV 21, we found that the hMN-ALS line had a 3.2-fold longer IBI (p < 0.0001) than the healthy hMN line (Figure 3c), while the hDN-PD line showed a 0.6-fold shorter IBI (p < 0.0001) than the healthy hDN line (Figure 3d).

**Figure 3.**
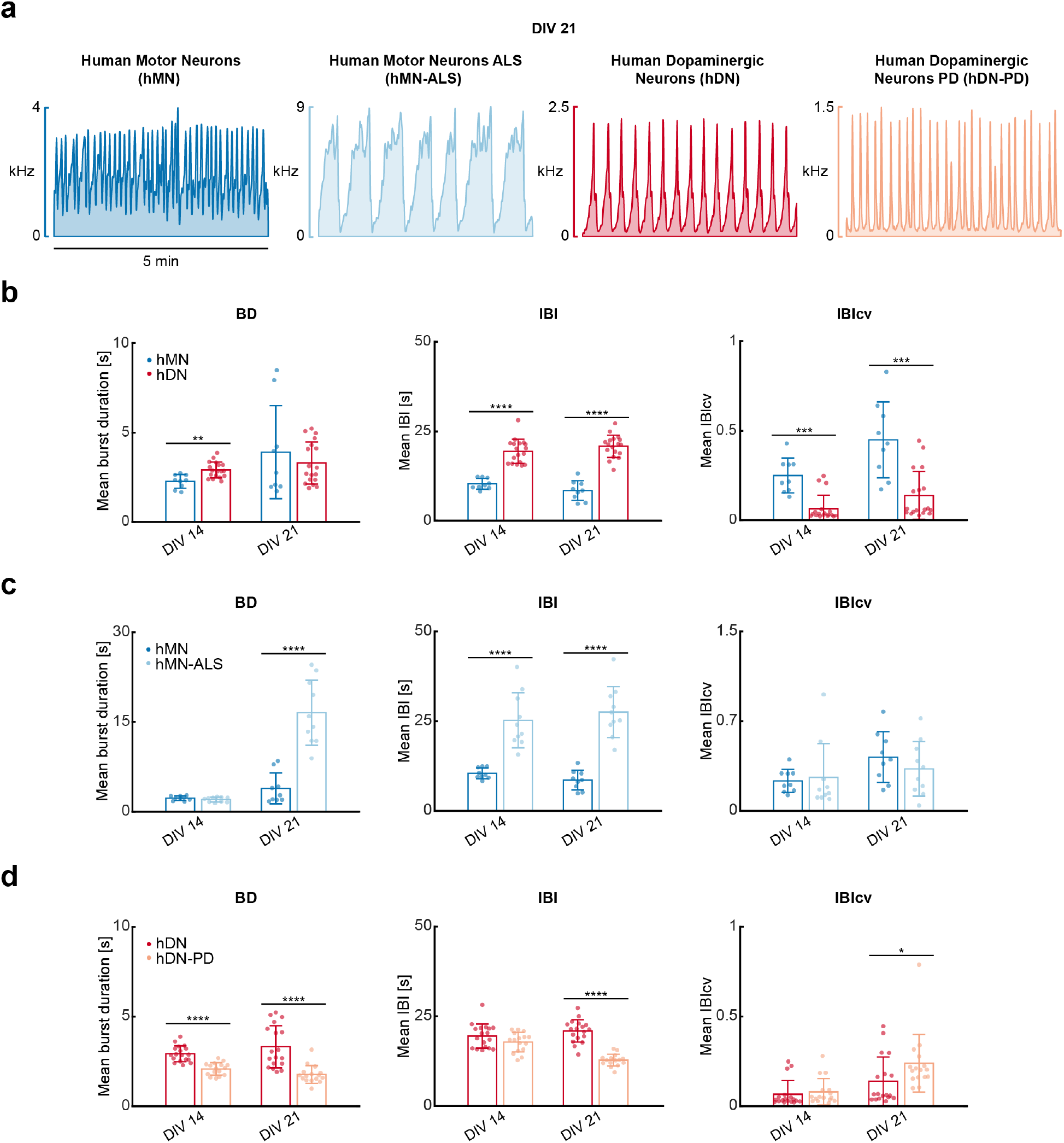
Network burst characterization of human iPSC-derived neurons across development. **(a)** Population spike time histograms simultaneously recorded by 1’024 electrodes from hMN (blue), hMN-ALS (light blue), hDN (red) and hDN-PD (orange) neurons at DIV 21. **(b)** Bar plots comparing mean burst duration (BD), mean inter-burst interval (IBI) and mean IBI coefficient of variation (IBIcv) of 9 hMN cultures (blue) and 18 hDN cultures (red) at DIVs 14 and DIV 21. Each dot represents one HD-MEA or well. **(c)** Bar plots comparing mean burst duration, mean IBI and mean IBI coefficient of variation of 9 hMN cultures (blue) and 10 hMN-ALS cultures (light blue) at DIV 14 and DIV 21. Each dot represents one HD-MEA or well. **(d)** Bar plots comparing mean burst duration, mean IBI and mean IBI coefficient of variation of 18 hDN cultures (red) and 16 hDN-PD cultures (orange) at DIV 14 and DIV 21. Each dot represents one HD-MEA or well. Bar heights indicate distribution mean values, and error bar indicate standard deviations. The black stars indicate p values: * p < 0.05, ** p < 0.01, *** p < 0.001, **** p < 0.0001.

An analysis of the IBI coefficients of variation yielded no significant differences between hMN and hMN-ALS lines (Figure 3c). However, the dopaminergic neuron lines showed a small difference at DIV 21 (p < 0.05) (Figure 3d).

### 2.4. Network-burst shape can be used to characterize human iPSC-derived neurons lines

By inspecting the population spike time histograms (Figure 4a), we noticed that each human iPSC-derived neuronal line featured a characteristic burst shape across every activity peak (see also Figure S2). Therefore, we assessed the ability to discriminate hMN and hMN-ALS or hDN and hDN-PD neurons by their network burst shape at a given day *in vitro*. For this purpose, we computed the linear correlations between every single recorded network burst and the corresponding *network burst templates* generated by averaging over many electrodes and HD-MEAs or wells harboring the same cell line at the specific DIV (see Methods, *HD-MEA Metrics*). In Figure 4b, we show the network burst template for hMNs at DIV 14 and DIV 21 (see Methods, *HD-MEA Metrics*). When we linearly correlated single recorded hMN bursts at DIV 14 to the DIV14 hMN template, we obtained an average Pearson Correlation Coefficient (PCC) of 0.96 ± 0.01, which indicated a high similarity between the burst template and the single bursts. However, upon correlating single DIV14 hMN-ALS bursts to the DIV14 hMN template, the average PCC decreased to 0.88 ± 0.06 (Figure 4b) and to 0.71 ± 0.12 for correlating DIV21 hMN-ALS bursts to the DIV21 hMN template (Figure 4b). In Figure 4d, we used the network burst templates of hDNs at DIVs 14 and 21. The DIV 14 hDN template yielded an average Pearson Correlation Coefficient (PCC) of 0.98 ± 0.01 upon application to single hDN recorded bursts at DIV 14. Upon using the DIV14 hDN template for single hDN-PD bursts recorded at DIV 14, the average Pearson Coefficient decreased to 0.96 ± 0.03 (p < 0.001) (Figure 4d). At DIV 21, the application of the DIV21 hDN template for DIV21 hDN-PD bursts yielded a further decreased PCC of 0.89 ± 0.03 (Figure 4d). In Figure 4c and 4e, the inverse correlations by using the hMN-ALS and hDN-PD disease line templates are shown. Expectedly, the correlations are stronger between the disease-line templates and the respective disease line bursts at both DIVs as compared to bursts of the healthy lines. As the disease lines featured a somewhat larger burst shape irregularity, the discrimination by using the disease line templates was less efficient in comparison to the healthy line templates.

**Figure 4.**
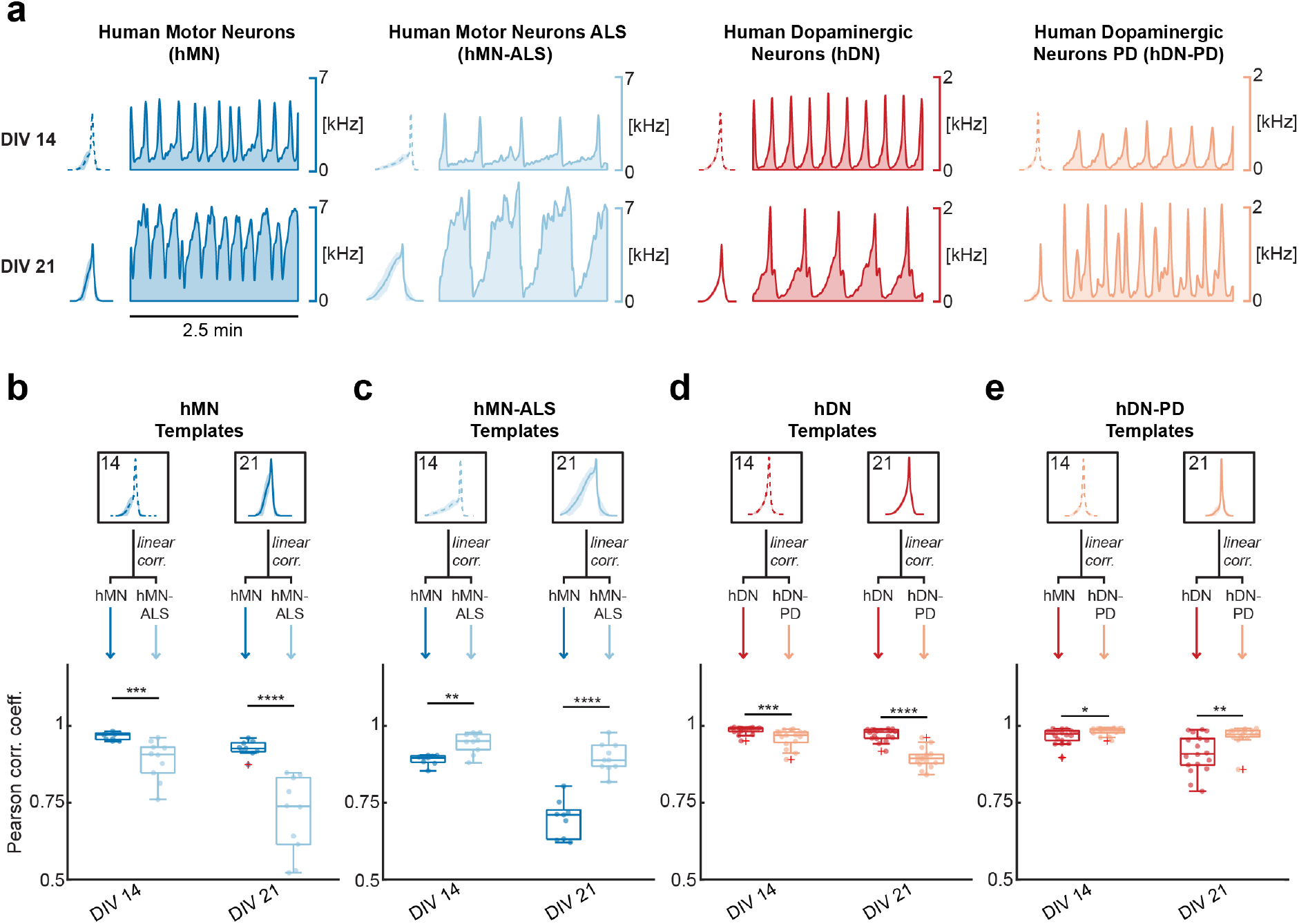
Network burst-shape characterization of human iPSC-derived neurons across development. (**a**) Population spike time histograms simultaneously recorded by 1’024 electrodes from (from left to right) hMN (blue), hMN-ALS (light blue), hDN (red), hDN-PD (orange) neurons at DIV 14 (top) and DIV 21 (bottom). At the left of each panel, an average network burst template is shown for the specific DIV and related neuronal cell type. **(b)** Bar plots comparing the Pearson Correlation Coefficient (PCC) upon linearly correlating the recorded network bursts to the corresponding average templates represented in the top panel. The graph represents the PCC upon linearly correlating burst of healthy motor neurons (N=9) and ALS motor neurons (N=10) to the template obtained from the healthy motor neurons at DIVs 14 and 21. Each dot represents one HD-MEA or well. Box plots indicate distribution mean value and standard deviation. **(c)** PCCs for correlating the hMN-ALS template to bursts of hMN (N=9) and hMN-ALS (N=10) lines. **(d)** PCCs for correlating the hDN template to burst of hDN (N=18) and hDN-PD (N=16) lines. **(e)** PCCs for correlating the hDN-PD template to bursts of hDN (N=18) and hDN-PD (N=16) lines. The black stars indicate p values: * p < 0.05, ** p < 0.01, *** p < 0.001, **** p < 0.0001.

### 2.5. Axonal action potential propagation in healthy and diseased human iPSC-derived neurons

The HD-MEA chip used in this work allows for reading out neuronal electrical activity at subcellular resolution^45–47^. Measurements of action potential propagation velocities and potential alterations can be used to functionally differentiate neuronal cell types^48^, to study axonal development^49,50^ and to get indications of axon degeneration^51^. Here, exploiting the high spatiotemporal resolution of HD-MEAs, we determined action-potential propagation velocities by tracking the AP propagation in space and time simultaneously across multiple electrodes (Figure 5a-5c and Figure S3). Action potentials are initiated at the AIS^46^, which produces the largest extracellular voltage signal^46^, and then propagate along the axon, while AP amplitudes decrease (Figure 5b).

**Figure 5.**
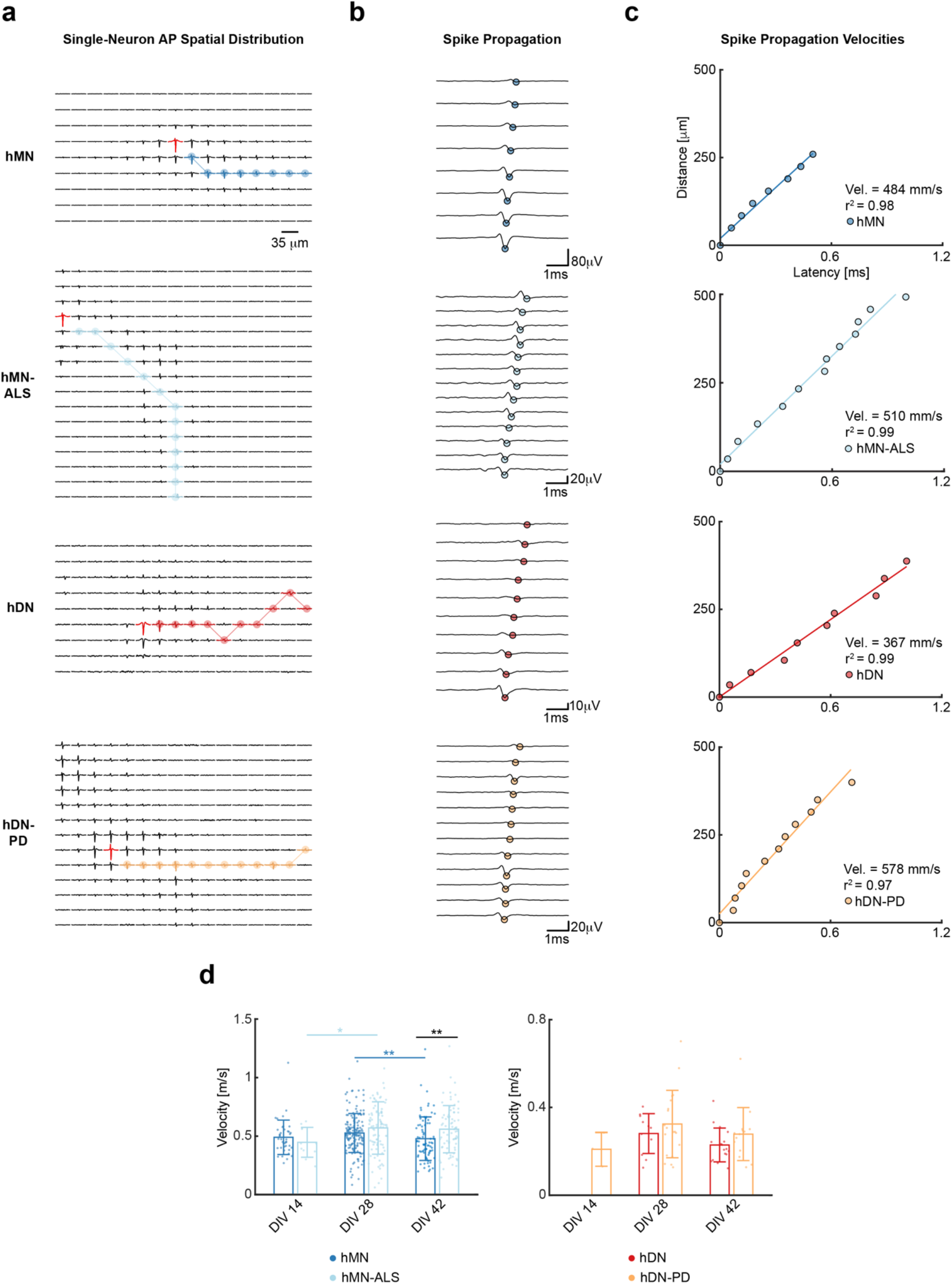
AP propagation velocity computed for different human iPSC-derived neuron lines. **(a)** Spatial distribution of action potential (AP) waveforms of sample neurons of (top to bottom) hMN, hMN-ALS, hDN, hDN-PD neuronal lines. Each trace represents a cutout of 6 ms of extracellular voltage signal recorded on the respective electrode at DIV 28. The red traces indicate the waveforms on the electrode featuring the largest signal amplitude. The plots are also displayed in Figure S3 at larger magnification. **(b)** AP propagation in time and space, after temporal alignment of the signals of selected readout electrodes represented in (a). Points indicate the voltage minima of the recorded traces. The traces are ordered with respect to delay from top to bottom with the top trace featuring the longest delay and coming from the electrode that is most distant to the axonal initial segment (AIS). **(c)** Linear regression interpolation to compute the AP propagation velocity for the neurons represented in (a) and (b). **(d)** Bar plots comparing the AP propagation velocity of (left) healthy motor neurons and ALS motor neurons and of (right) healthy and PD dopaminergic neurons, at DIVs 14, 28 and 42. Each dot represents one HD-MEA or well. Bar heights indicate distribution mean values, and error bars indicate standard deviations. Statistical significance is indicated using black stars to compare hMN and hMN-ALS neurons, blue stars to compare hMNs at different DIVs and light blue stars to compare hMN-ALS neurons at different DIVs. The stars indicate p values: * p < 0.05 and ** p < 0.01. The number (N) of hMNs for which velocities were determined at DIVs 14, 28 and 42 is 40, 165 and 80, respectively. The number (N) of hMN-ALSs for which velocities were determined at DIV 14, 28 and 42 is 11, 92 and 76, respectively. The number (N) of hDNs for which velocities were determined at DIVs 28 and 42 is 11 and 17, respectively. The number (N) of hDN-PDs for which velocities were determined at DIVs 14, 28 and 42 is 3, 19 and 14, respectively.

To test if axonal-action-potential-propagation velocities differed between healthy and diseased cell lines, we estimated (see Methods, *Action Potential Propagation Velocity)* the respective velocities of healthy motor neurons and ALS motor neurons and over several DIVs (14, 28 and 42). Results showed differences in velocities during more mature stages, i.e., at DIVs 28 and 42 (Figure 5d). At DIV 42, hMNs featured an average velocity of 440 ± 180 mm/s, while hMN-ALS neurons featured a significantly higher average axonal AP-propagation velocity of 560 ± 200 mm/s (p < 0.01). Moreover, the average hMN velocity decreased by 10% (p < 0.01) between DIV 28 and DIV 42, while the hMN-ALS average velocity increased by 30% from DIV 14 to 28 (p < 0.05).

The same measurements were conducted for healthy dopaminergic neurons and PD dopaminergic neurons. However, in contrast to motor neurons, healthy and diseased dopaminergic neurons did not show any significant difference in axonal propagation velocity (hDNs: 230 ± 80 mm/s, hDN-PDs: 280 ± 120 mm/s at DIV 42). The hDN lines featured comparably small signal amplitudes, which yielded less neurons that could be identified in the recordings. Although the spike sorting parameters were adjusted to also include neurons with smaller amplitudes (see Methods, *Spike Sorting)*, only few hDN neurons satisfied the propagation-velocity conditions described in Methods (*Action Potential Propagation Velocity*) and could be used to assess axonal propagation speeds.

### 2.6. Retigabine decreases spontaneous neuronal activity

Retigabine is an anticonvulsant drug used for epilepsy treatment. It opens neuronal K(v) 7.2-7.5 (formerly KCNQ2-5) voltage activated K(+) channels that generate the M-current, a subthreshold K(+) current needed to stabilize the membrane potential and control neuronal excitability^52^. The effects of retigabine dosage have been analyzed by monitoring neuron spiking rates using a 64-electrode MEA, which were found to significantly decrease^10,38,53^. Moreover, retigabine was also reported to decrease spontaneous electrical activity in neuronal cortical cultures by decreasing the mean burst rate^54^. Here, we explored the effects of retigabine on the inter-burst interval and number of active electrodes in cultures of healthy motor neurons at DIV14. Additionally, we investigated how the spiking rate estimation may be influenced by the number of readout electrodes.

An analysis over the entire HD-MEA electrode array showed a significant reduction in the fraction of active electrodes one minute after applying concentrations of 1 μM (p < 0.05), of 5 μM (p < 0.05) and 10 μM (p < 0.05) (Figure 6a, 6b).

**Figure 6.**
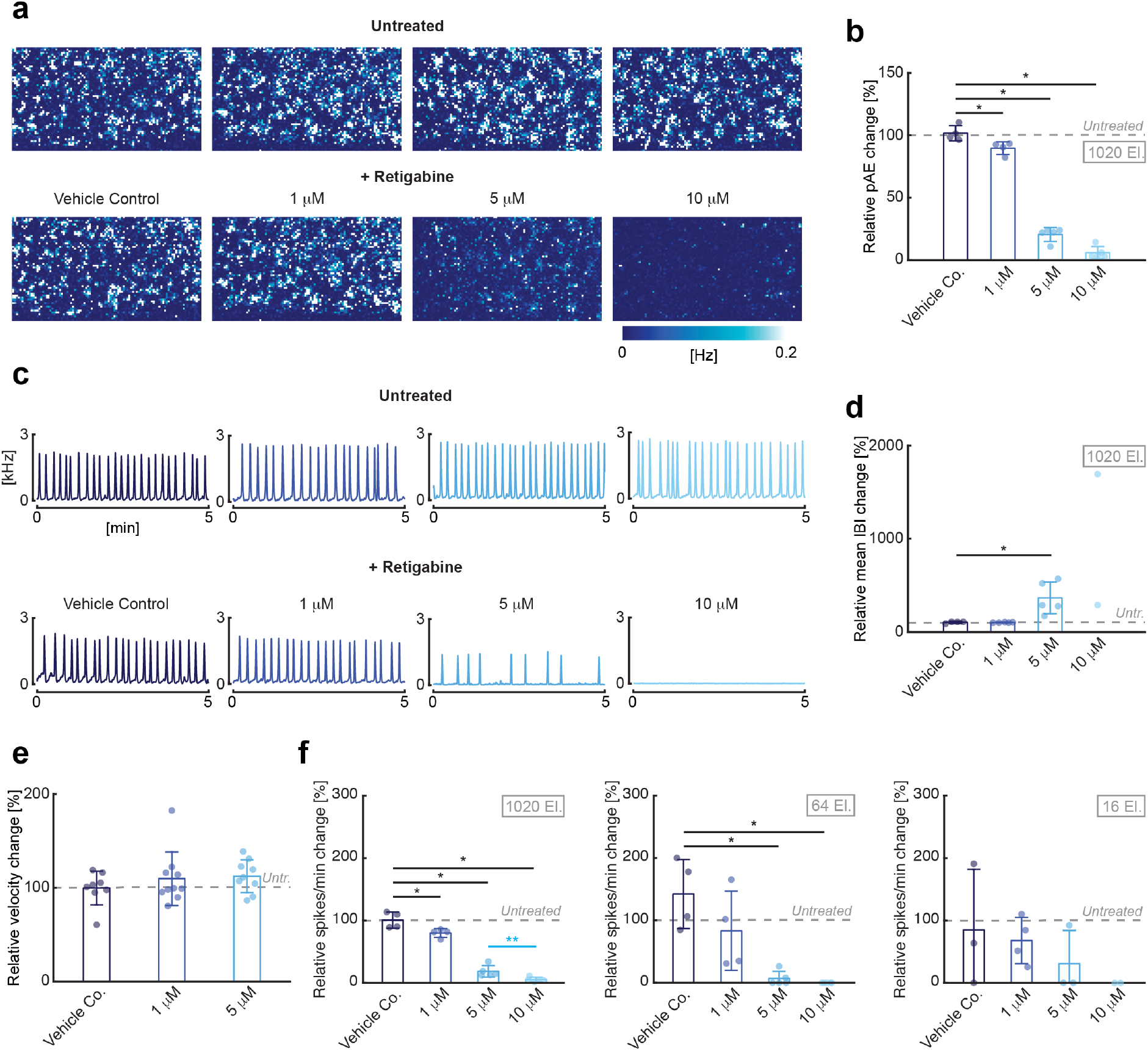
Retigabine effect on motor neurons on spikes/m, IBI and active electrodes. **(a)** 2D spatial distribution maps of electrode spike rates recorded from four exemplary HD-MEAs at DIV 14. Signals from 6’600 electrodes per HD-MEA were recorded before drug administration (top row) and one minute after drug administration (bottom row) for different retigabine concentrations. **(b)** Bar plots representing the relative change (percent) in active electrodes for each applied retigabine concentration and for the vehicle control with respect to pre-treatment conditions. Each dot represents one HD-MEA or well. Bar heights indicate distribution mean values and error bars indicate standard deviations. The dashed gray line marks the values (100%) before drug treatment. The number of HD-MEAs or wells included N=4 for vehicle control and N=5 for each retigabine concentration (1 μM, 5 μM and 10 μM). **(c)** Population spike time histograms of four representative HD-MEAs, recorded before (top row) and after drug administration (bottom row) for different retigabine concentrations. **(d)** Bar plots representing the relative change in the mean IBI for each applied retigabine concentration and for the vehicle control, normalized to pre-treatment conditions. **(e)** Bar plots representing the relative change in axonal velocity upon exposure to the vehicle control (N=8 neurons), 1 μM of retigabine (N=10 neurons) and 5 μM of retigabine (N=9 neurons), normalized to pre-treatment conditions. **(f)** Spikes/min-values computed as relative change with respect to the situation before drug administration. Plots show spikes/min-values for using signals from the 1’020 most active electrodes at a minimum pitch of 35 μm selected from the overall array of 26’400 electrodes (left), and configurations of 64 electrodes (center) and 16 electrodes at 200 μm pitch (right). For more details, see also Supplemental Figure 5a. The black stars indicate p values with respect to vehicle control. Light blue stars indicate p values between 5 μM and 10 μM retigabine concentrations. * p < 0.05, ** p < 0.01.

Using simultaneously 1’020 electrodes in the active regions, we assessed the number of spikes/min (see Methods, *Drug Administration)*, which was reduced to 80 ± 7% upon applying a concentration of 1 μM (p < 0.05), to 19 ± 9% upon applying concentrations of 5 μM (p < 0.05), and to 6 ± 4% using 10 μM (p < 0.05) (Figure 6f, left), compared to the baseline recordings without retigabine treatment. This activity reduction was expected, as retigabine hyperpolarizes the cell membrane by opening potassium channels.

We also determined the mean inter-burst intervals, which increased 3.5-fold (p < 0.05) upon administration of 5 μM of retigabine (Figure 6c, 6d). The inter-burst interval also increased upon applying 10 μM retigabine, albeit the effect was more difficult to assess as some HD-MEAs did not record any burst over the whole recording duration of 5 min.

Although retigabine caused significant changes in network firing properties and network bursting, retigabine doses of 1 μM and 5 μM did not significantly change the velocity of the propagating action potentials along axons (Figure 6e). Retigabine could potentially have altered AP propagation velocity along axons, as KCNQ channels are also present on dendrites and axon^55^. A very small decrease in conduction velocity was reported after retigabine administration in rat sciatic nerves^56,57^.

In a next step, we tested the influence of the number and configuration of used electrodes on the obtained spiking and IBI. In addition to the standard configuration comprising the 1’020 most active electrodes at a minimum pitch of 35 μm, we used configurations of 64 electrodes (grids of 8×8 electrodes at 200 μm pitch) and 16 electrodes (4×4 grid at 200 μm pitch), see also Figure S4a. By using the 64-electrode configuration, we found similar relative changes in spikes/min for concentrations of 5 μM (p < 0.05) and 10 μM (p < 0.05), however, we could not find differences between vehicle control and a concentration of 1 μM, and we noticed an increased variation of this metric across different cultures (Figure 6f, center). Using 16-electrode configurations, arranged in a 4×4 grid, we did not obtain significant differences between the control sample and retigabine dosage (Figure 6f, right), as the scattering of the measured values was comparably large. As can be expected, a larger number or recording electrodes or sampling points provides more reliable results.

To compare the variability in the measurement data and to see in how far local effects may influence the results, we applied 6 different configurations of 8×8 and 4×4 electrodes across 4 HD-MEAs for control and 1 μM concentration and 5 HD-MEAs for 5 μM and 10 μM concentrations (Figure S4b and 6d). For the metric spikes/min, we could confirm a higher well-to-well variability with respect to the configuration with 1’020 electrodes, while for the metric IBI, we found a significant difference between a drug concentration of 5 μM and no drug using the 64-electrode configuration, but we could not detect any effect of retigabine dosage using the 16-electrode configurations (Figure S4c).

## 3. Discussion and Conclusion

We presented an investigation of the electrical phenotypes of several human iPSC-derived neuronal cell lines using HD-MEAs. We showed that HD-MEA technology - despite the low signal amplitudes of the hiPSC-derived neurons of approx. 30 μV to 45 μV - enables a reliable discrimination of the electrical activity of different neuronal lines at different developmental stages and across several levels, ranging from whole-network activity levels to subcellular structures. As more and more human iPSC-derived neuronal lines become available^58^, the possibility to extract and compare a multiparametric set of physiological features across different functional levels will improve (i) the functional characterization of those neuronal lines in healthy and diseased states and (ii) the evaluation of drug effects upon administration *in vitro.*

As a first step, we characterized all cell lines by investigating metrics, such as mean firing rate (MFR), mean inter-spike interval (ISIcv), mean spike amplitude (MSA) and percentage of active electrodes (pAE) (Figure S1 and Figure 2). In a next step, we extracted features from network bursts of neuron cultures on MEAs, including burst structure, oscillatory behavior and synchronicity^59^ (Figure 3 and 4). Finally, we characterized neuronal lines by investigating axonal propagation velocities (Figure 5). In contrast to passive MEA devices that feature fixed electrode configurations, CMOS-based HD-MEAs offer the possibility to extract all aforementioned metrics from several active regions within a culture^23,60^. This feature enabled a reliable distinction of rPCNs from hDNs and hMNs based on the MFR, ISIcv, MSA, and pAE. (Figure S1 and Figure 2).

As for healthy and diseased motor neurons, hyperexcitability was reported in iPSC-derived motor neurons harboring A4V SOD1 mutations^10^ and C9ORF72 repeat expansions^61^. In studies with motor-neuron-like cell lines, transfected with mutant Q331K TDP-43, the authors found that the TDP-43 mutation increased the firing frequency of action potentials^62,63^. In our characterization of motor neuron electrical activity, we did not find a higher mean firing rate or mean spike amplitude in hMN-ALS neurons as compared to control hMNs (Figure 2). In contrast, we found that the hMN-ALS burst duration was 4.2-fold longer (16 sec) than that of control hMNs at DIV 21 (Figure 3).

As for healthy and diseased dopaminergic neurons, a higher network-bursting activity of A53T α-synuclein PD neurons, in comparison to a human control line, was reported by Zygogianni et al.^64^, while other studies of network properties of PD neurons with mutations in the LRRK2 gene found that diseased neurons lacked synchronous network bursting activity^65^. Here, we found an increased frequency in synchronized population bursting activity for PD neurons, which is in agreement with the findings of Zygogianni et al.^64^ (Figure 3). Furthermore, we found that PD neurons featured shorter burst duration and more variable inter-burst intervals compared to healthy dopaminergic neurons (Figure 3), which, again, is in agreement with the findings of Zygogianni ^64^.

Comparing the healthy and diseased lines by their AP propagation velocity, we found an increased axonal AP-conduction velocity in Q331K TDP-43 neurons (Figure 5), which could be correlated with altered axonal excitability properties reported in previous ALS studies^66,67^. Human motor-neuron lines feature larger AP spike amplitudes than dopaminergic-neuron lines, so that a more reliable extraction of axonal signals and differences in axonal propagation velocity between cell lines at different DIVs was possible. However, it is important to notice that the commercially available human iPSC-derived lines, used in this study, do not contain 100% motor neurons (≥ 75% for hMNs, see www.fujifilmcdi.com) or 100% dopaminergic neurons (≥ 80% for hDNs, see www.fujifilmcdi.com) (Figure 1), so that caution is advised in the interpretation of the data. For a reliable comparison of subcellular features of genetically defined cell lines it is necessary to make sure that only neurons of the specified genetic type are characterized, which can be achieved by using the following experimental techniques or combinations thereof: (1) purification of neuronal lines by FACS sorting and subsequent plating^10^; (2) activation and identification of genetically-defined neurons in a mixed population by expression of opto- and chemogenetic markers, driven by cell-type-specific promoters^68^; (3) identification of cell types by extraction and classification of extracellular AP waveform features^69,70^.

In a prototype drug dosage experiment, we evaluated the drug effects upon administration *in vitro* (Figure 6). We used retigabine, a potassium channel opener, which decreases neural activity. As expected, retigabine significantly decreased the percentage of active electrodes, the number of spikes per minute, and increased the inter-burst interval with increasing concentration (Figure 6). At the same time, we did not find significant changes in AP velocity across upon increasing drug dosage (Figure 6). We found that the number and density of used electrodes strongly influence the nature and number of metrics that can be used to assess drug effects (Figure 6). Additionally, our results showed that the use of a large number of flexible recording sites entailed a significantly lower intra-culture and culture-to-culture fluctuation as compared to applying fixed electrode configurations.

In conclusion, HD-MEA technology enables to conduct non-invasive assays and to extract highly reproducible metrics, such as mean firing rate, mean spike amplitude and burst shape, for studying disease mechanisms at a functional level for a multitude of human neuronal lines^71^. In addition, HD-MEA technology provides accurate readouts at single-cell and subcellular levels, which can be used as complementary metrics for assessment of drug effects in human neuronal cultures^72^.

## 4. Materials and Methods

### High-Density Microelectrode Arrays

CMOS-based HD-MEAs^23,60^ were used to record the extracellular signals of human iPSC-derived and rat primary neurons. The HD-MEA features 26’400 electrodes organized in a 120 × 220 grid within a total sensing area of 3.85 × 2.10 mm^2^. The electrode area is 9.3 × 5.45 μm^2^, and the center-to-center electrode distance (pitch) is 17.5 μm, which allows for recording of cell electrical activity at subcellular resolution. A user-selected configuration of 1’024 electrodes can be simultaneously recorded from (see also Methods, *HD-MEA Recordings*). The HD-MEA features noise values of 2.4 μVrms in the action potential band of 0.3 - 10 kHz and has a programmable gain of up to 78 dB. The sampling frequency is 20 kHz. We mostly used the MaxOne HD-MEA produced by MaxWell Biosystems AG (www.mxwbio.com).

### Cell Lines

Human iPSC-derived neurons and astrocytes (iCell® DopaNeurons, iCell® Motor Neurons, iCell® DopaNeurons A53T, iCell® Motor Neurons Q331K, iCell® Astrocytes) were purchased from FCDI (FUJIFILM Cellular Dynamics International, Madison, USA). We used two dopaminergic-neuron cell lines: a mixed population of healthy A9 and A10 subtype human dopaminergic neurons (hDNs), which have been demonstrated to express relevant midbrain dopamine neuronal markers (see also Figure 1), and an isogenic variant harboring the early-onset mutation A53T α-synuclein (hDN-PD) as a disease line. The A53T α-synuclein mutation renders α-synuclein susceptible to aggregation, which has been proposed as contributor to PD pathology^73^.

We used healthy human spinal motor neurons (hMNs) with expression of characteristic motor neuron markers (e.g. SMI-32), and an isogenic diseased motor neuron cell line carrying the TDP-43 Q331K mutation (hMN-ALS). Dominant mutations in the RNA/DNA-binding protein TDP-43 have been linked to amyotrophic lateral sclerosis (ALS)^74^. Human neuronal cell lines were co-cultured with human astrocytes at a ratio of 5:1. Rat primary neurons were obtained from dissociated cortices of Wistar rats at embryonic day 18, using the protocol described in Ronchi et al., 2019^75^. All animal experimental protocols were approved by the Basel-Stadt veterinary office according to Swiss federal laws on animal welfare and were carried out in accordance with the approved guidelines.

### Cell Plating

Prior to cell plating, HD-MEA chips were sterilized using 70% ethanol for 30 minutes. Ethanol was then removed, and the chips were rinsed three times with sterile tissue-culture-grade water. The HD-MEA chips were coated with chemicals to render the surface more hydrophilic. We adopted two different coating protocols suggested by the FCDI guidelines, depending on the cell type: for the motor-neuron plating, we treated the surface poly-D-lysine (PDL, 20 μL, 0.1 mg/mL) (A3890401, Gibco, ThermoFisher Scientific, Waltham, USA) for 1 hour at room temperature; for the dopaminergic-neuron plating, we used poly-L-ornithine (PLO, 20 μL, 0.05 mg/mL) solution (A-004-C, Sigma-Aldrich, St. Louis, USA) for 2 hours in a 5% CO2 cell culture incubator at 37 °C. The PDL and PLO solutions were then aspirated, and the electrode surfaces were rinsed three times with sterile water.

On the plating day, extracellular-matrix-protein solutions were added to promote cell adhesion. Before plating the motor neurons, Geltrex extracellular-matrix solution (10 μL, 0.16 mg/mL) (A1569601, Gibco) was pipetted onto the array and left for 1 hour at room temperature. Before plating the dopaminergic neurons, Laminin solution (10 μL, 80 μg/mL) (L2020-1MG, Sigma-Aldrich) was pipetted onto the array for 30 minutes at 37 °C.

Frozen neuron and astrocyte vials were thawed for 3 minutes at 37 °C in a water bath, thereafter, 7 mL of medium was added to dilute the dimethyl sulfoxide (DMSO). The vials were then centrifuged for 5 minutes at 1600 rpm. Neurons and astrocytes were resuspended in medium to the desired ratio of 100’000 neurons and 20’000 astrocytes and then plated onto the HD-MEA chips in a 10 μL medium drop. The cells on the HD-MEA chips were incubated in a 5% CO2 cell-culture incubator at 37°C for 1 hour. After cell adherence to the HD-MEA-chip surface, each well was filled with 0.6 mL of maintenance medium (see below). A 50% medium exchange was performed one day after plating, followed by 33% medium exchanges twice a week.

Maintenance medium consisted of Brainphys (05790, STEMCELL Technologies, Vancouver, Canada), supplemented with 2% iCell neural supplement (type B for dopaminergic neurons, type A for motor neurons) (FCDI), 1% iCell nervous-system supplement (FCDI), 1% of 100X N-2 supplement (17502048, Gibco), 0.1% laminin (L2020-1MG, Sigma-Aldrich) and 1% of 100X penicillin-streptomycin (15140122, Gibco).

During the first week *in vitro*, motor-neuron medium was supplemented with 5 μM DAPT (D5942, Sigma-Aldrich) to prevent outgrowth of proliferative cells.

E18 rat primary cortical neurons were isolated, plated and maintained according to protocols described previously^75^.

### HD-MEA Recordings

Recordings were performed weekly, starting from DIV 7. We used the “Activity Scan Assay” and “Network Assay” modules, featured in the MaxLab Live software (MaxWell Biosystems AG, Zurich, Switzerland), to monitor and record neuronal electrical activity. Seven electrode configurations, including a total of 6’600 electrodes at a pitch of 35 μm (every second electrode in x and y), were used to map spontaneous neuronal electrical activity across the entire HD-MEA chip. Each electrode configuration was recorded during 120 seconds (“Activity Scan Assay”). After scanning the entire HD-MEA, we selected 1’024 electrodes from the identified *active* electrodes and recorded network electrical activity for 300 seconds (“Network Assay”). Active electrodes were identified based on their firing rate, and among those the 1’024 electrodes featuring the highest firing rates were selected. The number of used HD-MEAs per condition (N) is always indicated in the figures in the Results section.

### Data Analysis

Data analysis was performed using custom-written codes in MATLAB R2019a and Python 3.6.10.

### HD-MEA Metrics

To characterize and compare the neuronal cultures, we used two categories of metrics: *electrode* metrics and *well* metrics. Metrics were first computed at electrode level. Then, the electrode metrics were averaged to represent a single HD-MEA chip or well. A *spike* was defined as a negative voltage deflection, whose amplitude exceeded 5 times the standard deviation of the baseline noise. Metrics indicated with asterisks (*) were adapted from Tukker et al., 2018^42^ and Cotterill et al., 2016^76^. The following parameters and metrics were used to assess neuronal characteristics.

1. *Electrode firing rate (eFR)* was defined as the number of spikes 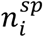 observed in a predefined time interval *T*, recorded on a single electrode *i*:

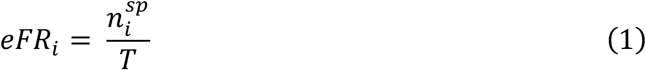
2. *Well mean firing rate* (MFR)* was defined as the mean of the eFR computed over the most active (electrodes with highest eFR) 1’024 electrodes (*N* = 1’024), simultaneously recorded:

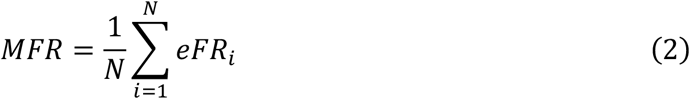
3. *Electrode spike amplitude (eSA)* was defined as the 90^th^ percentile of the recorded spike amplitudes on a single electrode *i*:

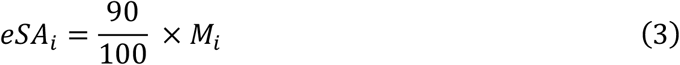

where *M_i_* is the length of the corresponding spike-amplitude vector, with amplitudes sorted from the smallest to the largest value.
4. *Well mean spike amplitude (MSA)* was defined as the mean of the eSA, computed over the most active 1’024 electrodes (*N* = 1’024), simultaneously recorded:

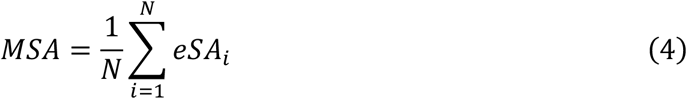
5. *Electrode inter-spike interval coefficient of variation (eISIcv)* was defined as the interspike interval (ISI) standard deviation, divided by the ISI mean (*μISI)*, where the ISI is the time difference between two consecutive spikes times *t_j_*, recorded on a single electrode *i*:

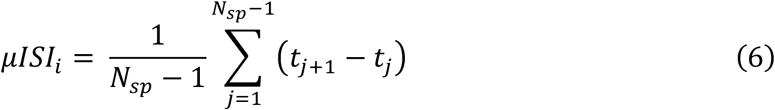

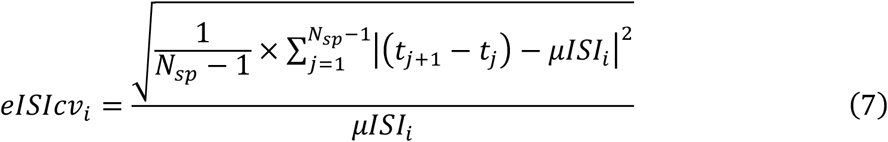
6. *Well inter-spike interval coefficient of variation* (ISIcv)* was defined as the mean of the *eISIcv*, computed over the most active 1’024 electrodes (*N* = 1’024), simultaneously recorded:

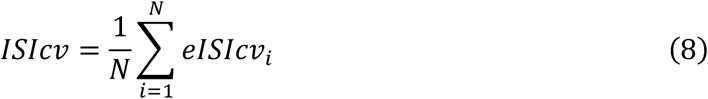
7. *Well percentage of active electrodes (pAE)* was defined as the percentage of electrodes with eFR > 0.1 Hz and eSA < −15 μV, per well. All electrode metrics described above (1-6) and all well metrics described below (8-11) were only computed for electrodes satisfying these two conditions for *active* electrodes (eFR > 0.1 Hz and eSA < −15 μV).
8. *Well mean burst duration* (BD)* was defined as the average duration of the network oscillations (bursts), recorded in a well:

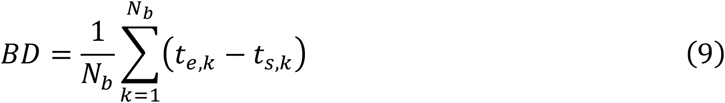

where *N_b_* is the number of bursts, *t_e_* represents the burst end point and *t_s_* the burst starting point as illustrated in Figure S5. The fast Fourier transform (FFT) was used to compute the main oscillation frequency and to correct for false positives as a consequence of high oscillation frequencies within each burst.
9. *Well mean inter-burst interval* (IBI)* was defined as the average time between the starting points of consecutive network bursts per well:

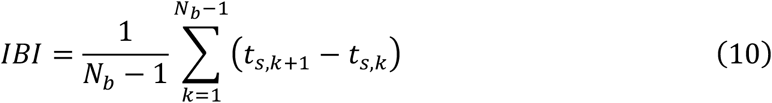
10. *Well mean inter-burst interval coefficient of variation* (IBIcv)* was defined as the inter-burst interval standard deviation, divided by the inter-burst interval mean (*IBI)* per well:

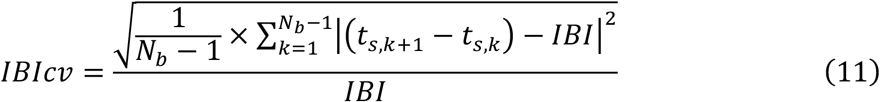
11. *Well Pearson correlation coefficient (PCC)* was defined as the average linear correlation between the network bursts of a well and a network burst template representing a specific cell type:

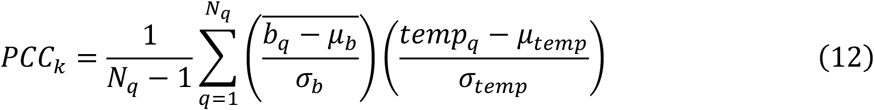

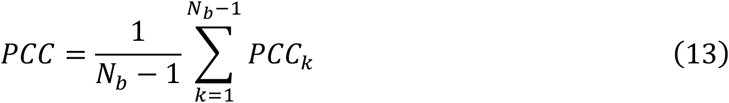

where *PCC_k_* represents the PCC between a single burst *b* (of mean *μ_b_* and standard deviation *σ_b_*) and a template *temp, N_q_* the number of observations in the burst and *PCC* the mean of all *PCC_k_* values computed in a specific well.

To characterize cell types according to their burst shape, eight templates were created, covering all cell types (hMN, hMN-ALS, hDN, hDN-PD) and each recorded DIV (14 and 21). The templates (temp) were obtained by averaging all recorded bursts in a well (*temp_w_*) and across wells of the same cell type and DIV^77^:

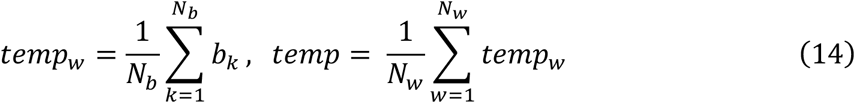

where *N_b_* is the number of bursts per well, *b_k_* denotes a single burst in a well, and *N_w_* is the total number of wells featuring a specific cell type and DIV.

### Spike Sorting

Spike sorting was performed to identify single units in the extracellular recordings. We ran spike sorting on the recordings, with which we covered the entire chip area (“Activity Scan Assay”) by using seven configurations of 960 electrodes (35 μm electrode pitch). We used the Kilosort2^78^ software within the SpikeInterface^79^ framework and the corresponding default parameters. Due to the large number of recordings, we automatically curated the spike sorting output according to the following criteria: for motor neurons, we removed clusters with less than 100 spikes, an ISI violation rate^80^ above 0.3, and a signal-to-noise ratio (SNR) below 5; for dopaminergic neurons featuring less activity and lower signal amplitudes (Figure 3b), we removed clusters with less than 50 spikes and an SNR below 3 (the threshold on ISI violation rate was the same). Automatic curation was performed using the SpikeInterface package^79^. Action potential propagation velocity

Spike sorted and curated units were used to estimate the propagation velocity of action potentials. We discarded units with a *small spatial extension* template (averaged extracellular waveforms), and only kept units whose template covered at least 10 electrodes with an amplitude of more 10% of the maximum amplitude of the given unit. With these templates, we used a graph-based approach to find distinct neuronal branches. For each template, we considered the electrodes featuring a signal with an amplitude of at least 5% of the maximum amplitude and with a kurtosis value above 0.3 to filter out channels without a distinct signal peak. We then sorted the selected electrodes according to the time difference of the signal peak occurrence on that electrode with respect to the signal peak occurrence on the electrode showing the maximum signal amplitude, which most likely was close to the axon initial segment^46^. We then built a graph with the selected electrodes as *nodes* (Figure S6). Starting from the nodes with latest peak occurrence times (largest time difference)^81^, each node was connected with *edges* to the three nearest electrodes with an earlier peak time. These three electrodes then formed the next set of nodes from which the procedure continued. Once the graph with all edges and nodes was built, we looked for the shortest paths along the nodes from each node to the electrode where the initial signal occurred, again starting from the electrodes with latest peak times or largest time differences. If an electrode formed already part of a path, it could not be used for another path. Finally, we removed duplicate paths by discarding the paths, where 50% of the nodes were in close proximity (<50 μm) to nodes of other paths. For the selected branches, distances were computed as the cumulative distance of all nodes between the initial electrode and the end of the path, while the peak times at the nodes included the differences between the peak time at the respective electrode and that at the initial electrode. Velocities were then estimated using a linear regression on the peak time differences and cumulative distances. We discarded branches with an *r^2^* <0.9, and in cases that the algorithm found more than one branch for a template, we kept only the one with the highest *r^2^*.

### Statistical Analysis

The statistical analysis was performed using the non-parametric Wilcoxon rank sum test for Figure 2 to 6. Given two populations, the null hypothesis assumes that it is likely that a randomly selected value from one population will be different from a randomly selected value from the other population. The test was used for the small sample sizes in Figure 2, 3, 4, 6 (N < 20 HD-MEAs for each cell line) and for the non-normal distribution of the data in Figure 5. For Figure 6, the Wilcoxon rank sum test was used to compare the multiple values obtained for drug treatments with the value for the vehicle control.

### Drug Administration

Retigabine (90221, Sigma-Aldrich), dissolved in DMSO (D2650, Sigma-Aldrich) was applied at concentrations of 1 μM, 5 μM and 10 μM to a co-culture of healthy human motor neurons and astrocytes, similar to the experimental procedures reported by Wainger et al., 2014^10^. The vehicle control consisted of culture medium and DMSO (D2650, Sigma-Aldrich).

We performed recordings of healthy motor neurons on each HD-MEA immediately before retigabine administration at DIV 14. We then conducted measurements one minute after drug administration.

Drug effects were quantified about one minute after dosage by using the following metrics: spikes/min (*S_/min_*), percentage of active electrodes (*pAE*) and mean inter-burst intervals (*IBIs*) (see Methods, *HD-MEA Metrics*).

The metric *spikes/min* (Figure 6b) for 1’024, 64 and 16 electrodes was computed as:

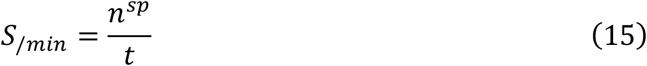

Where *n^sp^* is the total number of spikes recorded from all the recording electrodes during the recording time t in minutes. The resulting metric value, obtained from a drug-treated HD-MEA chip, was then normalized to the corresponding metric value, obtained from an untreated HD-MEA chip, in order to obtain a relative metric change (percent change) induced by the addition of the compound.

### Microscopy and Staining

Neurons on HD-MEAs were fixed using a 4% paraformaldehyde solution (FB001, ThermoFisher Scientific). Fixation prevented cell necrosis and autolysis and preserved the cellular constituents. Samples were then permeabilized and blocked using a BPS 10× (AM9625, ThermoFisher Scientific) solution containing 10% normal donkey serum (NDS) (017-000-001, Jackson ImmunoResearch, West Grove, USA), 1% bovine serum albumin (BSA) (05482, Sigma-Aldrich), 0.02% Na-Az (S2002, Sigma-Aldrich) and 0.5% Triton X (93443, Sigma-Aldrich). Permeabilization facilitated antigen access to the cell, while blocking prevented non-specific binding of antibodies to the tissue. Primary and secondary antibodies were diluted in a PBS solution containing 3% NDS, 1% bovine serum albumin (BSA), 0.02% Na-Az and 0.5% Triton X^75^. The used antibodies are listed in Table 1.

**Table 1.** Antibodies. The table describes primary and secondary antibodies used in this work.

| Antibody | Supplier | Ratio | Binds to | Catalogue # |
| --- | --- | --- | --- | --- |
| Anti MAP2 | EnCor | 1:500 | Microtubules | CPCA-MAP2 |
| Anti TH | Pel Freeze | 1:1000 | Tyrosine hydroxylase enzyme | P60101-150 |
| Anti GFAP | Abcam | 1:500 | GFAP | Ab7260 |
| Anti S-100 | Abcam | 1:200 | Zn | Ab52642 |
| Anti Smi-32 | Abcam | 1:500 | Neurofilaments | Ab7795 |
| Hoechst | Thermofisher Scientific | 1:500 | DNA | H3570 |

We imaged cells on the HD-MEA chip with a Nikon NiE upright confocal microscope, with a Yokogawa W1 spinning-disk scan head.

## Acknowledgements

This work was supported by the European Community through the European Research Council Advanced Grant 694829 “neuroXscales” and the corresponding proof-of-concept Grant 875609 “HD-Neu-Screen”, the Swiss Project CTI-No. 25933.2 PFLS-LS “Multi-well electrophysiology platform for high-throughput cell-based assays”, and through the Swiss National Science Foundation under contract 205320_188910 / 1. This work was also supported by an ETH Zürich Postdoctoral Fellowship 19-2 FEL-17 to Alessio Paolo Buccino. The funders had no role in study design, data collection and analysis, decision to publish, or manuscript preparation.

We thank Peter Rimpf and Tomislav Rebac for post-processing CMOS-based HD-MEAs. We thank the BEL group at ETH Zürich for valuable scientific discussions throughout the project. We thank Urs Frey, Marie Obien and David Jäckel at MaxWell Biosystems for their contributions to data analysis and interpretation. We thank Xinyue Yuan (ETH Zürich) for input to the axon-propagation-velocity computation. We thank Sam Malkin for proofreading the manuscript.

## Author contributions

S.R. contributed to the work design, performed and planned the experiments, analyzed and interpreted data, wrote the manuscript. A.P.B contributed to spike sorting data and velocity computation. G.P. contributed to perform experiments and to velocity computation. S.S.K. contributed to writing the manuscript. M.S. contributed to writing the manuscript. M.F. coordinated and conceived the project, contributed to work design and data interpretation and wrote the manuscript. A.H. coordinated and conceived the project and contributed to writing the manuscript. All the authors have approved the submitted version and revised it.

## Competing financial interests

The authors S.R., A.P.B., G.P, S.S.K., M.S. and A.H. declare no competing financial interests. M.F. is co-founder of MaxWell Biosystems AG, which commercializes HD-MEA technology.

