## Supplementary Material for "Electrophysiological Phenotype Characterization of Human iPSC-Derived Neuronal Cell Lines by Means of High-Density Microelectrode Arrays"

**Figure S1. Characterization of rat primary cortical neurons across development**

To have a comparison for the electrical activity patterns of the human iPSC-derived neuronal lines, the developmental stage of which may greatly vary, and for the metrics described in the Methods section, we measured the electrical activity of rat primary cortical neurons (rPCNs), the most commonly used neuronal cells in the microelectrode-array field and an established *in-vitro* culturing system, across development. We plated rPCNs on 5 HD-MEAs, and we followed their development from DIV 7 to DIV 21.

The electrical activity of the rPCNs was assessed by recording spontaneous neuronal action potentials across the whole HD-MEA-chip active area and by then computing the following metrics: (1) mean firing rate (MFR), (2) mean spike amplitude (MSA), (3) mean inter-spike interval (ISI) coefficient of variation (ISIcv) and (4) percentage of active electrodes (pAE) (see Figure S1a, S1b and Methods, *HD-MEA Metrics*). rPCN cultures showed a 3.8-fold increase in the mean firing rate ($p < 0.01$, Wilcoxon rank sum test) from DIV 7 (0.54 ± 0.07 Hz) to DIV 21 (2.07 ± 0.99 Hz), as well as a 5.6-fold increase in the percentage of active electrodes ($p < 0.01$) from DIV 7 (8.6 ± 1.7 %) to DIV 21 (48.6 ± 15.7 %) (Figure S1c). The mean spike amplitude increased 1.5 fold ($p < 0.01$) from DIV 7 (64.9 ± 7.6 μV) to DIV 14 (95.7 ± 7.7 μV) and then stabilized between DIV 14 and DIV 21 (95.0 ± 9.0 μV) (Figure S1c). The mean ISI coefficient of variation increased 2.16 fold ($p < 0.05$) during culture development, with values of 1.53 ± 0.08 at DIV 7 and 3.31 ± 0.26 at DIV 21, which indicated that the firing rate in rat neurons became less regular during development (Figure S1c). Overall, rPCNs showed an increase in mean firing rate and percentage of active electrodes across development, as well as an increase in mean spike amplitude and irregularity of neuronal spiking (Figure S1c), as has been reported in other studies characterizing the electrical activity of neocortex neurons by MEA technology[44,78].

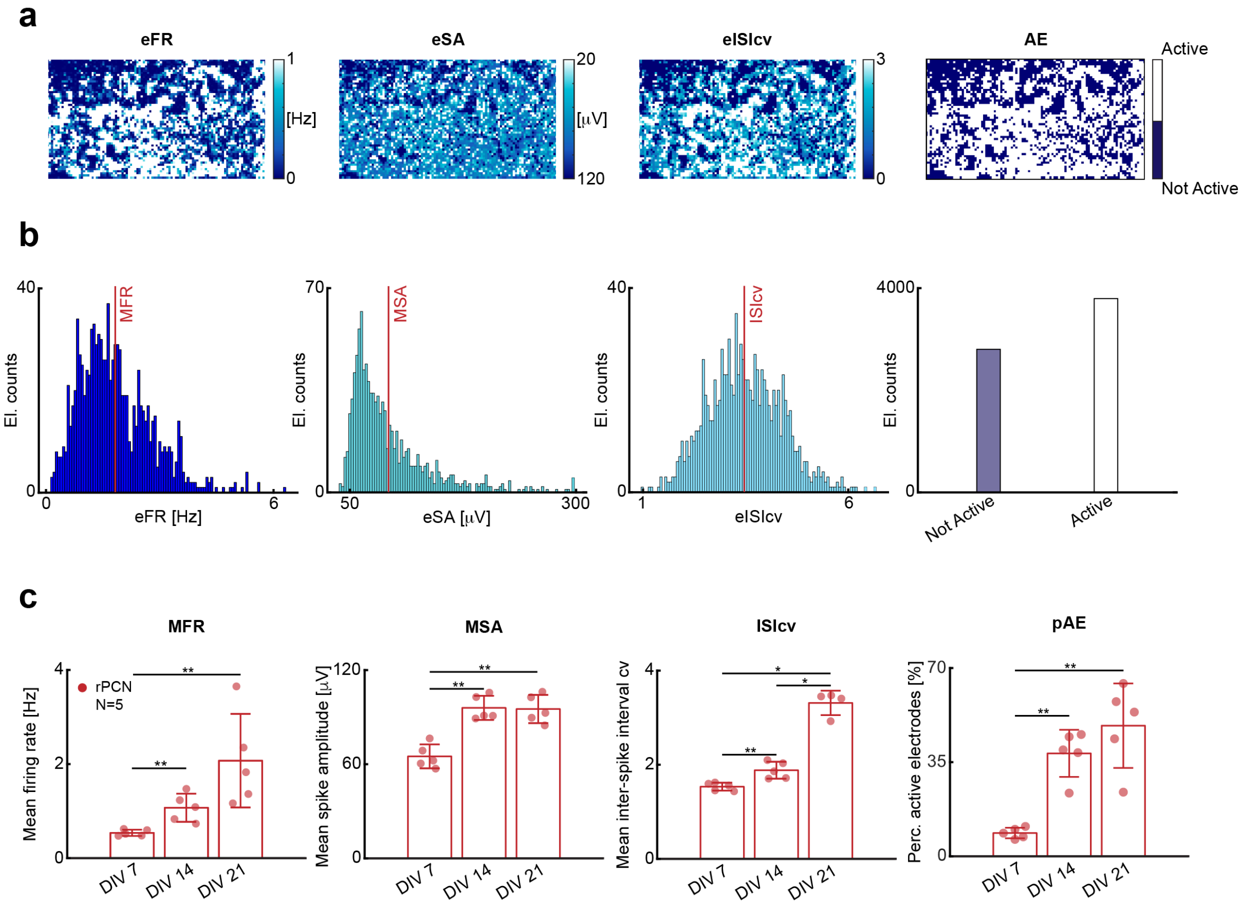


**Figure S1. Electrical phenotype characterization of rat primary cortical neurons across development.**

**(a)** Exemplary 2D spatial distribution maps of electrode firing rates (eFR), electrode spike amplitudes (eSA), electrode ISI coefficients of variation (eISIcv) and active electrodes across a total of 6'600 electrodes (see Methods, *HD-MEA Metrics*). The colors in each map represent the measured value of the given metric. The four maps have been acquired from one rPCN HD-MEA chip at DIV 21. **(b)** Exemplary distributions of metrics for one rPCN HD-MEA chip at DIV 21: first panel: distribution of eFRs; second panel, distribution of eSAs; third panel, distribution of eISIcvs; fourth panel, distribution of pAEs. Distribution means are indicated by red vertical lines. **(c)** Bar plots comparing mean firing rate (MFR), mean spike amplitude (MSA), mean ISI coefficient of variation (ISIcv) and percentage of active electrodes (pAE) of 5 different rPCN HD-MEAs (N = 5) at DIVs 7, 14 and 21 (see Methods, *HD-MEA Metrics*). Each red dot represents one HD-MEA or one well. The bar heights indicate the distribution mean values, and error bars indicate standard deviations. The black stars indicate p values: * $p < 0.05$, ** $p < 0.01$.

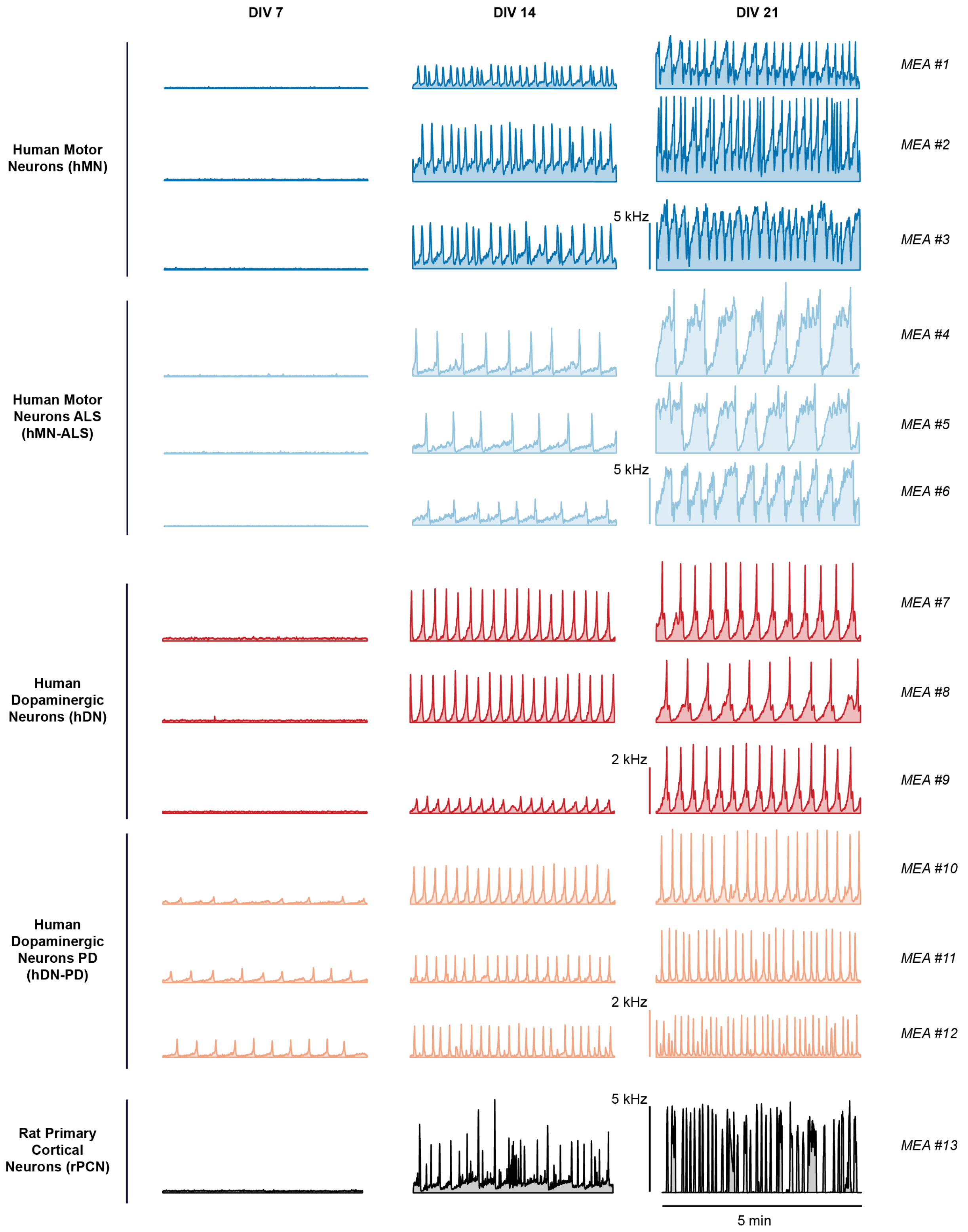


**Figure S2. Burst reproducibility across samples and HD-MEAs.**
Population spike time histograms simultaneously recorded from 1’024 electrodes of hMN (blue), hMN-ALS (light blue), hDN (red), hDN-PD (orange) and rPCN (black) cultures at DIVs 7 (first column), 14 (second column), 21 (third column). Bars at the left of the graphs in the third column indicate the spiking frequency.

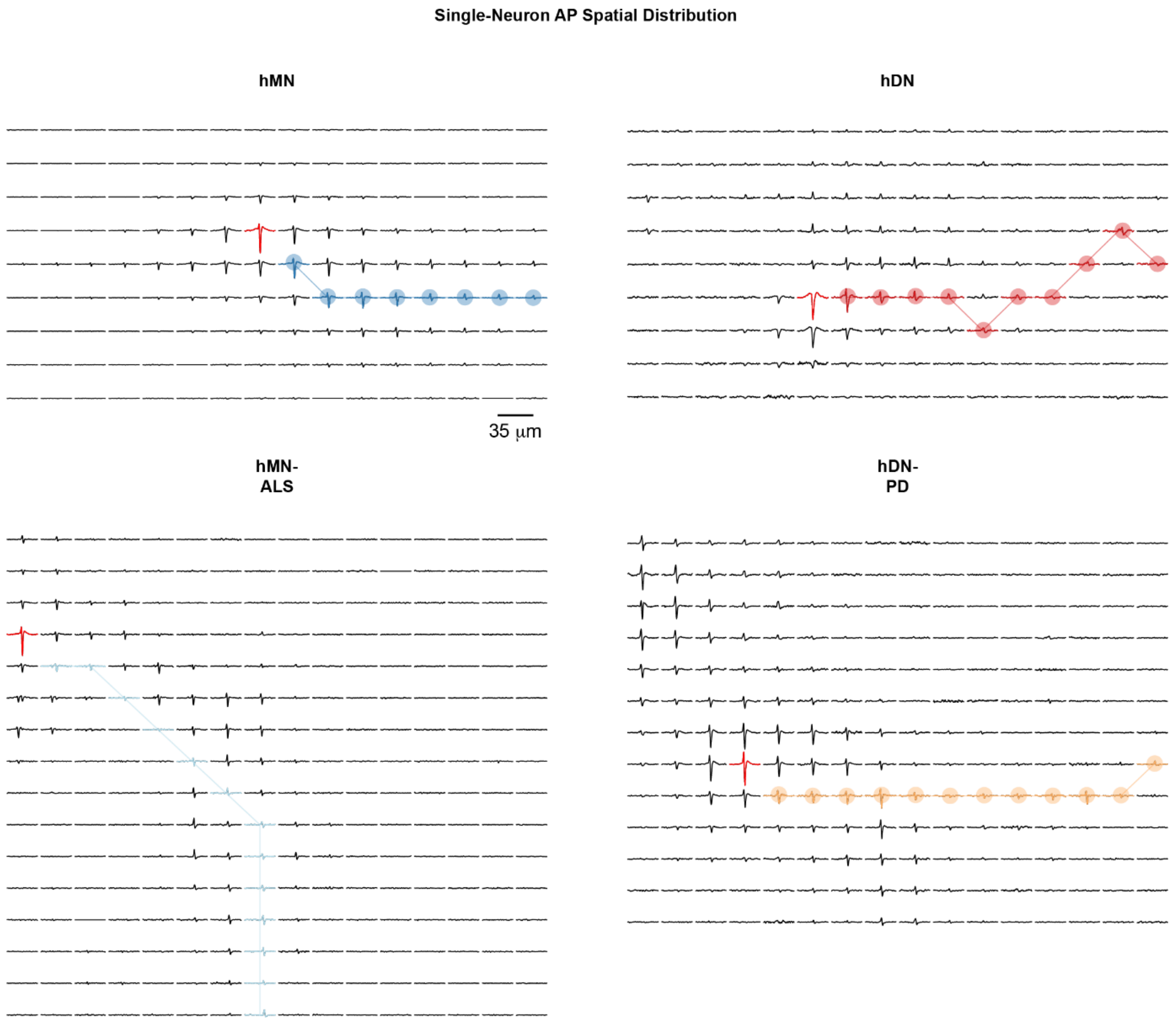


**Figure S3. Single-neuron action potential spatial distributions.**
Spatial distribution of action potential (AP) waveforms of sample neurons of hMN (top left), hMN-ALS (bottom left), hDN (top right), hDN-PD (bottom right) neuronal lines. Each trace represents a cutout of 6 ms of extracellular voltage signal, recorded on the respective electrode at DIV 28. The red traces indicate the waveforms on the electrode featuring the largest signal amplitude. The plots are identical with those in Figure 6a and are enlarged for better visibility.

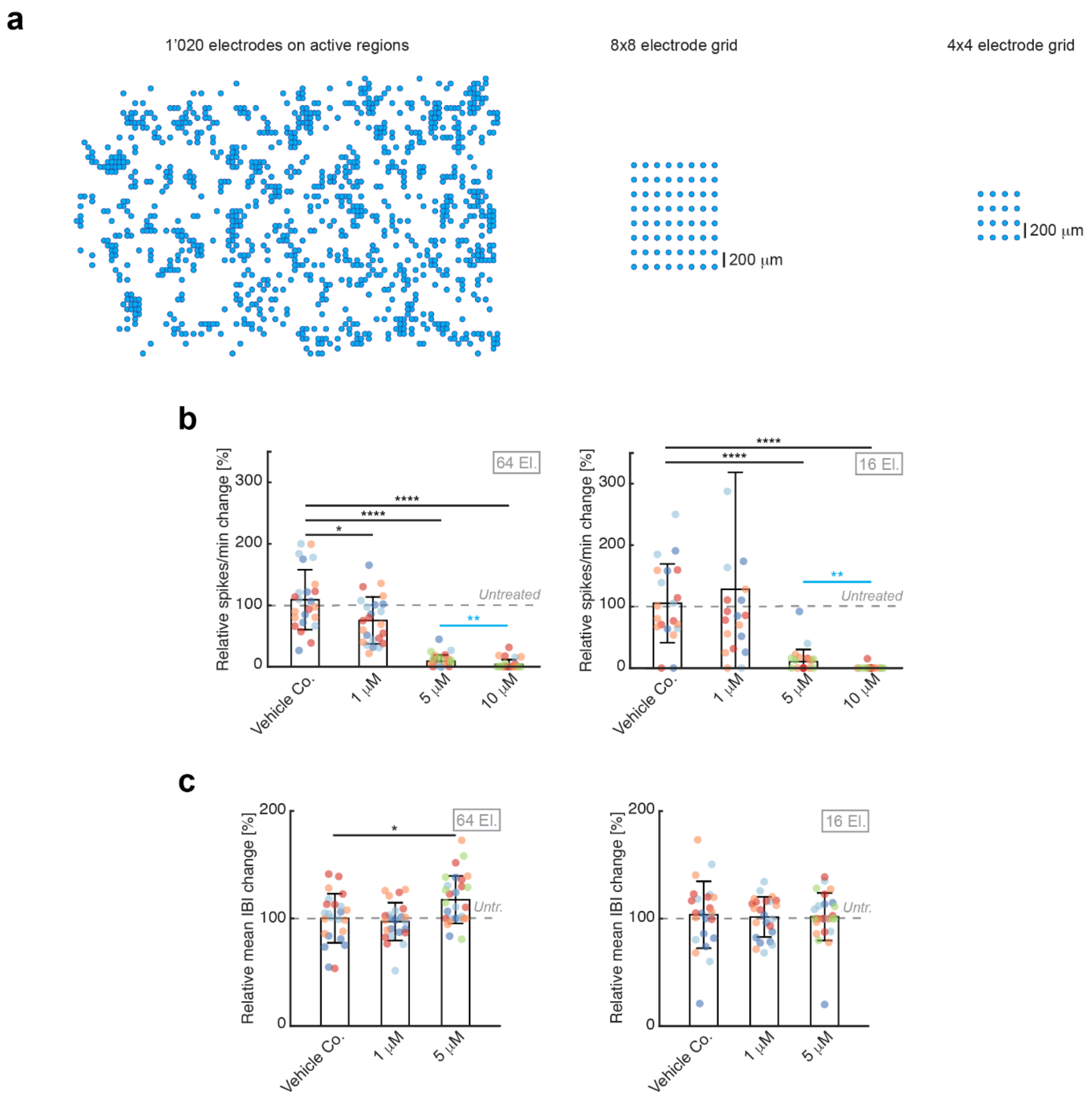


**Figure S4. Retigabine well-to-well variability as a function of recording electrodes.** **(a)** Electrode configurations to assess retigabine effects described in Figure 7b and 7f. For the first configuration (left), 1'020 electrodes at a minimum pitch of 35 µm were selected from the active regions across the whole HD-MEA, i.e., the 1'020 electrodes recording the highest firing rates. We also used 8×8 (center) and 4×4 (right) electrode grid configurations with 200 µm electrode pitch. Each electrode configuration was measured before and after retigabine dosage to assess the drug effects. **(b)** Six different configurations of 8×8 and 4×4 electrodes over 18 HD-MEAs (N=4 for vehicle control and 1 µM concentration, N=5 for 5 µM and 10 µM concentrations) were used to assess variability of the results obtained with the fixed electrode configurations across a culture with respect to variations from culture to culture and to see in how far local effects may influence the results obtained with those low-density configurations. The measurement points originating from low-density electrode configurations in the same culture are indicated in the same color. Bar plots represent the relative change in spikes/min upon exposure to vehicle control, 1 µM, 5 µM and 10 µM of retigabine, normalized to pre-treatment conditions. Bars indicate distribution mean values, and error bars indicate standard deviations. The dashed gray line marks the values (100%) before drug treatment. **(c)** Six different configurations of 8×8 and 4×4 electrodes over 13 HD-MEAs (N=4 for vehicle control and 1 µM concentration, N=5 for 5 µM concentration) were used to assess variability across a culture with respect to variations from culture to culture and to see in how far local effects may influence the results obtained with those low-density configurations. The measurement points originating from low-density electrode configurations in the same culture are indicated in the same color. Bar plots represent the relative change in mean IBI upon exposure to the vehicle control, 1 µM and 5 µM of retigabine, normalized to pre-treatment conditions. The black stars indicate p values with respect to vehicle control. Light blue stars indicate p values between 5 µM and 10 µM retigabine concentrations. * $p < 0.05$, ** $p < 0.01$, **** $p < 0.0001$.

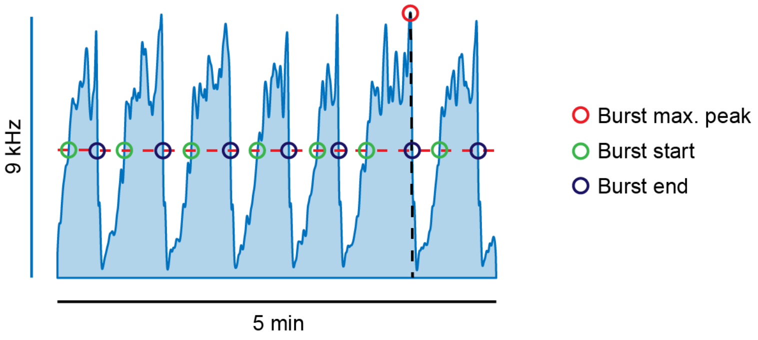


**Figure S5. Burst duration computation.**
Representative spike time histogram recorded by 1’020 electrodes. The burst duration was computed by fixing a threshold (dashed line) at 50% of the maximum burst amplitude (red). Start (green) and end points (blue) were placed at the intersection of spike time histogram and threshold.

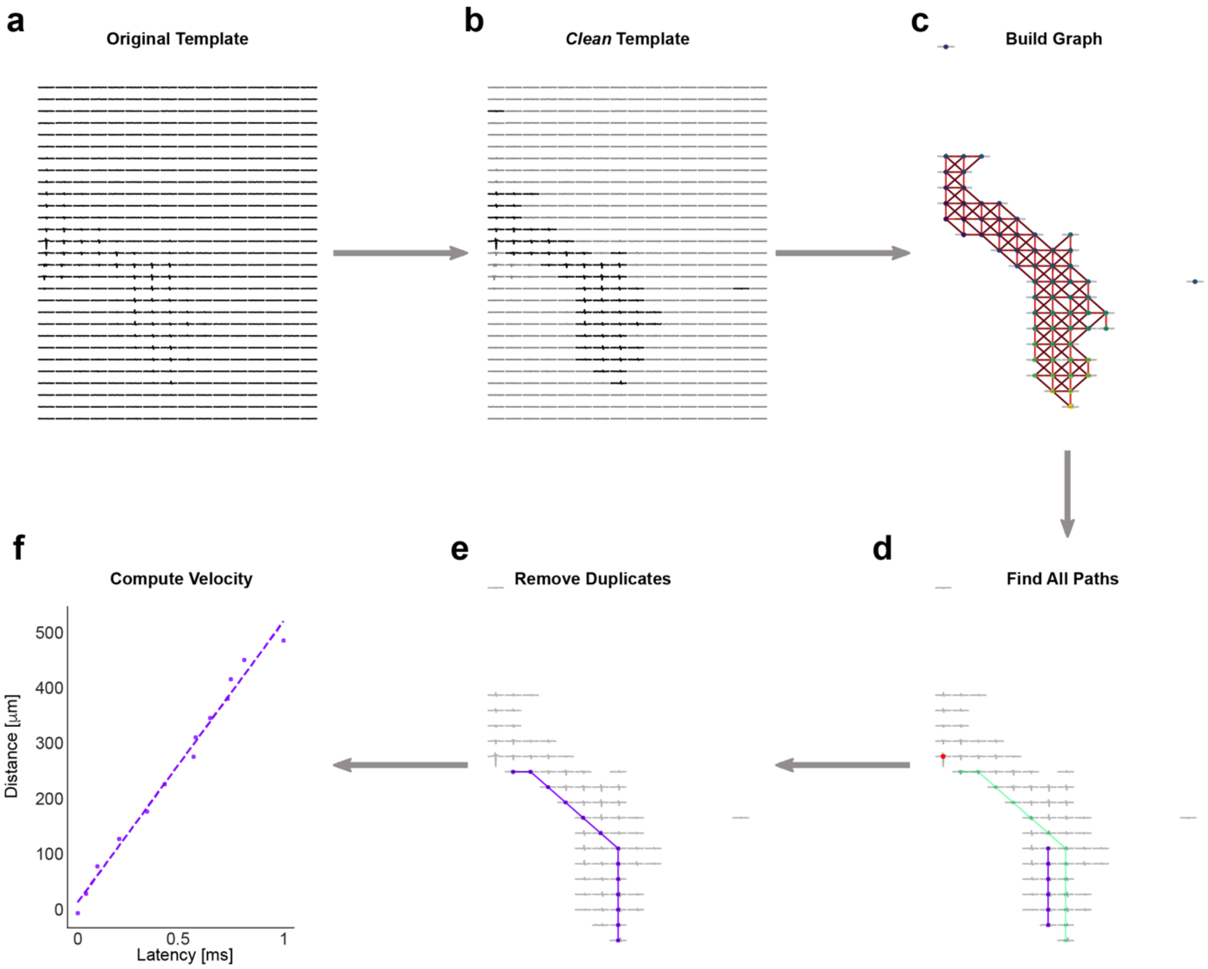


**Figure S6. Axonal propagation velocity algorithm**.
**(a)** Original template of a neuron obtained after spike sorting. **(b)** Selection of a subset of electrodes to obtain a "clean" template based on 3 criteria: (i) spike amplitude on a selected electrode > 5% of the maximum signal amplitude of the respective neuron; (ii) the signal kurtosis on a selected electrode > 0.3 (in order to filter out electrodes without a clear peak); and (iii) the peak time point of a selected electrode must occur after the peak time of the electrode with the highest amplitude (*initial electrode*). **(c)** The nodes of the graph correspond to the electrodes selected according to the procedure in (b). Starting from the nodes with latest peak occurrence times (largest time difference), each node was connected with edges to the three nearest electrodes with an earlier peak time within a distance of 50 µm. These three electrodes then formed the next set of nodes from which the procedure continued. If an electrode formed already part of a path, it could not be used for another path **(d-e)**. Duplicate paths removal by discarding the paths, where 50% of the nodes were in close proximity (<50 µm) to nodes of other paths. **(f)** Velocity estimation using a linear regression on the peak time differences and cumulative distances. Branches with an $r^2 < 0.9$ were discarded, and in cases that the algorithm found more than one branch for a template, only the one with the highest $r^2$ was kept.
